# Ecological interactions mediate evolutionary responses to temperature in microbial communities

**DOI:** 10.64898/2026.02.13.705791

**Authors:** Ewaldo Leitão, Megan Liu, Andrea Yammine, Ze-Yi Han, Katrina DeWitt, Douglas Chalker, Jean P. Gibert

## Abstract

Microbial populations play a pivotal role in ecosystem-level responses to rising temperatures and both their ecology and evolution can be directly influenced by warming. However, predicting microbial evolution and its ecological consequences is challenging because different genotypes within a population might respond uniquely to shifts in the abiotic and biotic environment. To understand how, we quantified evolutionary and ecological responses across temperatures in a protist of wide geographic distribution in the presence and absence of other microbial species with whom they interact (i.e., heterospecifics). In the absence of heterospecifics, we found that intraspecific interactions and warming selected in favor of a particular genotype, reducing genotypic diversity. In the presence of heterospecifics, 1) genetic diversity was further reduced under warming, resulting in temperature-dependent selection; but, 2) the magnitude of this change depended on the sign (+, 0, -) of the net ecological effect on the focal species by the heterospecifics, and this effect was itself temperature-dependent. Together, our results demonstrate that both intra- and interspecific interactions can mediate how temperature shapes microbial population rapid evolutionary responses, underscoring the importance of the ecological context in predicting evolutionary outcomes under climate change.

## Introduction

A central challenge in ecological research is to understand how species interactions may unfold in a rapidly changing climate (Åkesson et al., 2021; Blois et al., 2013; Gilman et al., 2010; Post, 2013), particularly when species evolve within ecological timescales (Dam et al., 2021; Hairston Jr et al., 1999; O’Donnell et al., 2018; Pantel et al., 2015; Urban et al., 2016). This rapid evolution is shaped by ecological interactions within communities, which modulate the strength and direction of selection driving evolutionary change (Pimentel, 1968, 1961; Salamin et al., 2010; Tseng and O’Connor, 2015). In turn, rapid evolution alters the ecological interactions that impose selection and shape phenotypes, which is often referred to as the eco-evolutionary feedback (De Meester et al., 2019; McPeek, 2017; Vanvelk et al., 2024; Yoshida et al., 2003). These eco-evolutionary feedbacks are commonplace, but they are especially likely in microbial communities, where ecological and evolutionary timescales overlap strongly (Govaert et al., 2021; Han et al., 2023; Vanvelk et al., 2024).

Microbes are ubiquitous and vital to ecosystem functioning, providing important ecosystem-level functions that ultimately fuel the flux of matter and energy within and across ecosystems (Azam et al., 1983; Lindeman, 1942; Wetzel, 1995). Their large population sizes, high genotypic diversity, and short generation times, make them ideal to study community-level evolutionary dynamics in ecological timescales (Collins et al., 2020; Ghoul and Mitri, 2016). Understanding eco-evolutionary processes in microbial communities is therefore critical for predicting how climate change will alter ecosystems, yet it is a mostly unaddressed issue in climate change ecology (Barbour and Gibert, 2021).

Rising temperature influences organismal physiology (Bernhardt et al., 2018; Dell et al., 2011), often resulting in thermal performance curves (or thermal responses), which describe how growth rates and other performance traits vary across temperatures (Stark et al., 2025; Wieczynski et al., 2021). These physiological responses can affect the strength of ecological interactions (O’Connor, 2009; Vasseur and McCann, 2005), reshaping community- and ecosystem-level dynamics (Bonnaffé et al., 2024; O’Connor et al., 2009; Yvon-Durocher et al., 2010).

Increasing temperatures can drive thermal adaptation (Barneche et al., 2021; Barton et al., 2020; Liu et al., 2024; Padfield et al., 2016) and environmental change rewires species interactions, which in turn reshape the selection landscape imposed by those interactions (Barbour and Gibert, 2021; Bartley et al., 2019; Wieczynski et al., 2026). However, how both processes may simultaneously determine evolutionary responses to rising temperatures, is not known (Han et al., 2024). Such biotic shifts are often overlooked in studies of thermal adaptation, even though they alter selection regimes (Barbour and Gibert, 2021; Carroll et al., 2023).

A more fundamental understanding of these eco-evolutionary consequences under climate change therefore hinges on studying species-specific evolutionary responses along with the changes in the community context in which they occur (Govaert et al., 2021). Indeed, as the biotic and abiotic environment change, shifting ecological interactions should result in changes in relative fitness among genotypes in novel environments. While new genetic variants arise from mutation, recombination, or gene flow (Hendry et al., 2018; Lenormand, 2002), selection can and often does operate on standing genetic variation (Barrett and Schluter, 2008; Burke et al., 2014; Lai et al., 2019), resulting in rapid adaptive change (Barrett and Schluter, 2008; De Meester et al., 2019; Dobzhansky, 1950; Lyberger and Schoener, 2023). Such evolutionary change can in turn influences ecological dynamics: genotypes with distinct traits like competitive ability, growth rates, or resistance to predation, will differentially shape interspecific interactions within the broader community (Bolnick et al., 2011; Crutsinger et al., 2008; Han et al., 2024; Leitão et al., 2024; Noto and Hughes, 2020; Taylor et al., 2019; Wieczynski et al., 2026). Thus, an increased genotypic diversity could promote population performance through complementarity effects (Barbour and Gibert, 2021; Ellers et al., 2011; Fridley and Grime, 2010; Kotowska et al., 2010; Vellend, 2006), which may prevent extinctions and dampen impacts on the community (Bell and Gonzalez, 2009; Gomulkiewicz and Holt, 1995; Hermann and Becks, 2022). At the same time, these trait-mediated eco-evolutionary interactions are also shaped by heterospecific trait variation (Colom and Baucom, 2020; terHorst, 2011). Thus, genotypic diversity is both affected by environmental change –biotic or abiotic–, and a driver of ecological interactions, creating an eco-evolutionary feedback loop where evolutionary change under warming reshapes the very ecological context in which selection occurs.

While genotypic diversity and environmental change jointly shape eco-evolutionary feedbacks, the role of central ecological processes – intra- and interspecific interactions – in mediating responses to warming, remains unclear. Within a population, individual genotypes might vary in the thermal performance curve (TPC) of their intrinsic growth rates (*r*), or r-TPCs (Liu et al., 2024; Wieczynski et al., 2026). Because intrinsic growth rates are a measure of absolute fitness (Abrams et al., 1993; Lande, 1976), systematic changes in r with temperature among genotypes –i.e., change in fitness with temperature across genotypes– should result in temperature-dependent selection (Liu et al., 2024), and changes in relative genotypic frequency and diversity across temperatures (Fig 1A). However, genotype-genotype interactions may alter genotype-specific thermal responses, modifying population evolution (i.e., *intraspecific effects*, Fig 1B). Additionally, the presence of a heterospecific (i.e., *interspecific effects*) can either decrease genotype performance without changing the overall genotype-specific thermal response (Fig 1C), or fundamentally alter the genotype-specific thermal response, and thereby shift the evolutionary trajectory of the focal population (Fig 1D). Within this eco-evolutionary framework, interaction type is expected to generate distinct evolutionary outcomes. Intraspecific interactions may amplify temperature-dependent fitness differences among genotypes, strengthening selection and accelerating genotypic sorting. Meanwhile, interspecific interactions could reduce (e.g., in competition) or increase performance (e.g., in a mutualism) across genotypes, limiting evolutionary change despite ecological effects, or alter the shape of genotype-specific TPCs altogether, thus redirecting evolutionary trajectories.

**Figure 1:**
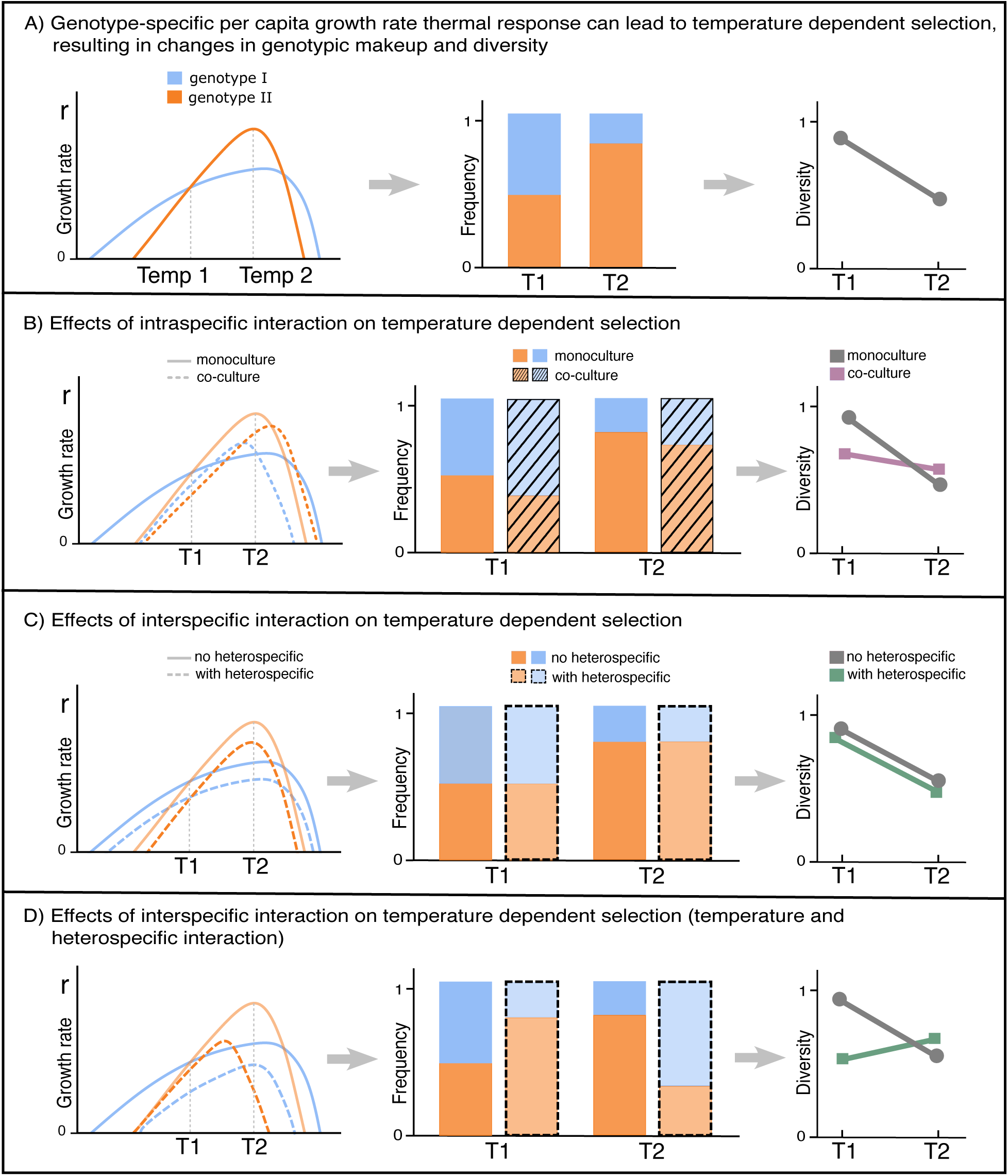
Conceptual illustration of the hypotheses tested. Panels depict alternative hypothetical scenarios of how temperature and biotic interactions could influence genotype-specific thermal performance curve of intrinsic growth rate (rTPC) and, in turn, genotypic composition. For each scenario, the panel shows the r-TPCs for each genotype (left), the corresponding expected population genotypic frequency (center), and genotypic diversity (right) at two discrete temperatures. A) Expected genotypic frequency and diversity based solely on individual genotypes thermal responses; here thermal responses are not changed by interspecific interactions. B) Intraspecific interaction (co-culture) changes genotype thermal responses, which alters genotypic frequency and diversity. C) Interspecific interaction (with heterospecifics) decreases the performance of the genotypes, without changing the thermal response, which does not alter the genotypic frequency and diversity. D) Interspecific interaction (with heterospecifics) decreases performance and changes the shape of the thermal response of genotypes, which results in changes in genotypic frequency and diversity.

Here, we investigated how temperature and intra- and inter-specific ecological interactions jointly affect the evolutionary and ecological responses of a microbial species. Specifically, we address the following questions: 1) can we predict change in genotypic frequency and diversity (i.e., evolution) based on the r-TPC of individual genotypes (i.e., intraspecific interactions)? 2) how does the presence of other heterospecifics (i.e., interspecific interactions) influence genotypic frequency and diversity? 3) do heterospecific traits (i.e., growth rate) play a role in it all?

To investigate these questions, we assessed whether and how populations of the protist *Tetrahymena thermophila* (the focal species) evolved differentially across temperatures through selection on standing variation (Barrett and Schluter, 2008; Lai et al., 2019) with or without three different heterospecifics; the protists *Colpidium striatum*, *Paramecium aurelia*, and *Paramecium bursaria*. All three species are widely assumed to be competitors of *Tetrahymena thermophila* (Carrara et al., 2015; Fox and Morin, 2001). Our results reveal evolutionary and ecological effects that are temperature dependent: decreasing genotypic diversity at warmer temperatures and clear effects of both intra- and interspecific interactions in these observed evolutionary shifts in genotypic frequency across temperatures. These effects are explained by a combination of differences in growth rates among heterospecific species, as well as their net effect (positive, neutral, negative) on the focal species’ population density.

## Methods

### Protist cultures

We focused on the protist *Tetrahymena thermophila*—a freshwater species that is distributed across the eastern United States (Zufall et al., 2013) and that is part of a genus of cosmopolitan distribution and importance (Lynn and Doerder, 2012). We used three protist species as heterospecifics: *Paramecium aurelia*, *Paramecium bursaria*, and *Colpidium striatum*. All these protist species are bacterivorous ciliates commonly found in aquatic ecosystems and have been widely treated as competitors of *Tetrahymena spp.* in the literature (Carrara et al., 2015; Fox, 2002; Fox and Morin, 2001). We characterized shifts in intraspecific genotypic variation in *T. thermophila* by keeping track of the frequencies of three distinct genotypes (IMB6, AXS, and DMCK72H), sourced from the Chalker lab (Washington University, Table S1). This simplified protist community, consisting of a focal species with multiple genotypes and a heterospecific species, represents a minimal community context sufficient to capture intra and interspecific interactions and their impacts on evolutionary dynamics. Upon reception of *T. thermophila* strains, we transferred all cultures from axenic Proteose Peptone growth medium to modified Timothy Hay growth medium (Altermatt et al., 2015) made with autoclaved dionized water and 2g/L of commercial Timothy Hay. The media was inoculated with a bacterial community from Duke Forest Gate 9 pond/Wilbur pond (Lat 36.013914, Long -78.979720), and an autoclaved wheat kernel as a carbon source (Altermatt et al., 2015). The bacterial community was fully described elsewhere and includes thousands of bacterial species (DeWitt et al., 2025; Rocca et al., 2022). Once we account for the complexity of the bacterial community, our microbial communities (protists + bacteria) are much more similar to what might be observed in nature than what is commonly used in the laboratory (Altermatt et al., 2015), and arguably represents an entire complex microbial community, despite its simplicity at the protist level. All cultures were maintained in Percival (Perry, IA) AL-22 growth chambers under light (12:12hr day:night cycle) and temperature controlled conditions (22°C) in 250mL borosilicate jars filled with 150mL of growth medium. Because *T. thermophila* is a natural bacterivore, these experimental conditions are more realistic than purely axenic ones.

### Growth rate responses to temperature

We quantified intrinsic growth rates of all protist species (and all *T. thermophila* genotypes) in 3cm diameter Petri dish microcosms with 3mL of Timothy Hay growth medium at three temperatures (19, 22, 25°C), each replicated six times. These temperatures capture the range most commonly experienced in their natural habitat during the growth season (Zufall et al., 2013). Microcosms were initialized by pipetting three individual cells from stock cultures under a scope (Leica stereomicroscope model M205C) and allowing them to grow for 24hrs, after which we censused the microcosms through whole-population counts under the scope. We calculated intrinsic growth rate, *r*, as ln(final density/initial density)/time (Wieczynski et al., 2021), with time = 1 day.

### Intra- and interspecific interaction assays

We operationally define ecological interactions as the net effect of the heterospecific on the population of the focal species, *T. thermophila*. These net effects can arise from direct effects (e.g., exploitative or interference competition) and/or resource-mediated indirect effects resulting from changes in the bacterial community the protists feed on. Together they demonstrate the cumulative net ecological effect of heterospecifics on the focal species’ density.

We set up multi-species assays with all three *T. thermophila* genotypes across the same three temperature treatments (19, 22, 25°C) in the presence of each of the other three protists (*P. aurelia*, *P. bursaria*, *C. striatum*), and a control without heterospecifics. All microcosms were replicated six times in a fully factorial design (temperature x treatment, n = 72). Microcosms were initialized at equal densities (5 individuals/mL of each *T. thermophila* genotype) in 6cm diameter plastic Petri dishes. Microcosms contained a total volume of 10 mL of Timothy Hay growth medium, and a wheat kernel. In treatments that included heterospecifics, microcosms were initialized to include additional 30 individuals/mL of each heterospecific. Experiments lasted 7 days, we sampled all microcosms to estimate genotypic frequencies at the end of the experiment.

Each *T. thermophila* genotype carries a different fluorophore gene (AXS – Yellow Fluorescent Protein, DMCK72H – mCherry fluorophore, IMB6 – Green Fluorescent Protein), that can be induced to fluoresce through addition of Cadmium Chloride (CdCl_2_) solution. We induced fluorescence using 0.1μL of a 1mg/mL CdCl_2_ solution per 100 μL of sampled volume per microcosm, for a final concentration of 1 μg/μL (Liu et al., 2024). Fluorescently tagged genotypes may lose their ability to fluoresce over time. Because of this, they all carry a Paromomycin resistance gene, so that fluorescing individuals can be selected for through Paromomycin exposure. We treated the microcosms with 100μg/mL of Paromomycin 24 hr before sampling, and 30 minutes after adding the CdCl_2_ we destructively harvested the entire microcosm for censusing. There is no record of the fluorescent marker influencing the fitness of the tagged strains, and our treatment comparisons are made relative to controls. Therefore, any possible marker effect would be accounted for in the baseline control. We used a Novocyte 2000R flow cytometer (Agilent, Santa Clara, CA) to count individual *T. thermophila* cells and estimate relative frequencies of each *T. thermophila* genotype based on fluorescence in 30 μL samples. Other protists were not counted at the end of the experiment.

### Statistical Analyses

To compare whether there are differences among *T. thermophila* genotypes’ r-TPC and genotypic frequencies across temperatures (without heterospecifics), we performed linear models with growth rate as the dependent variable and the interaction of genotype, as factor, and temperature, as a continuous covariate, as independent variables (growth rate ∼ genotype * temperature). We used count data for all genotypes to calculate the genotypic diversity of *T. thermophila* population, using the Shannon Diversity index 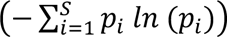, where *P_i_* is the proportion –or frequency– of individuals of each genotype *i* relative to total population size of *T. thermophila,* and *S* is the total number of genotypes (n = 3). While the Shannon Diversity index captures both richness and evenness of the genotypes, here it mostly captures changes in evenness because no genotype was lost or gained (genotypic richness is constant). These steps – calculation of genotypes’ r-TPCs in isolation, and genotypic frequencies and diversity when together– are those referred to in Figure 1.

To test for the effect of intraspecific interactions on genotypic diversity (without heterospecifics), we performed an ANOVA with temperature as the independent variable and genotypic diversity as the dependent variable. We tested for homoscedasticity and normality of the data and confirmed the model assumptions were met. Additionally, we used the linear model results of r-TPC of genotypes to generate expectations for genotypic frequency and diversity in the absence of intraspecific interactions.

To test for the effect of interspecific interactions on *T. thermophila* genotypic frequency and diversity across temperatures, we used linear models where frequency/diversity were dependent variables, and temperature as a covariate. For genotypic frequency, we modeled both genotype and heterospecific species identity as independent variables (frequency ∼ temperature * genotype * heterospecific). For genotypic diversity, heterospecific identity was used as the independent variable (diversity ∼ temperature * heterospecific).

To understand the mechanisms through which heterospecifics influenced evolutionary responses in *T. thermophila* across temperatures, we evaluated the effect of interspecific interactions on the final density of *T. thermophila* by calculating the effect size of the presence of each heterospecific species in the abundance of *T. thermophila*. This was done by comparing the controls (without heterospecifics) with the heterospecific treatments (i.e., effect size = *density*_treatment_ – *mean_density*_control_) at each temperature and for each heterospecific. We did so for each *T. thermophila* genotype separately, and for the total density of all genotypes in the population with log_10_ transformed data. This is similar to the approach used in (Jiang and Morin, 2007) – *mean_treatment – mean_control* –, but here we incorporated error from both treatment and control using bootstrap replicates (n = 999) to calculate 95% confidence intervals. We used effect size and confidence interval to infer neutral, negative, or positive effects of heterospecifics on *T. thermophila* density. If confidence intervals overlapped with zero, there was no effect of the heterospecific on *T. thermophila* population abundance. Effect sizes below or above zero, likely represent negative and positive effects of heterospecifics on *T. thermophila* abundance, and were therefore interpreted as such. We used the function ‘mean_diff’, from R package ‘dabest’ (Ho et al., 2019).

To evaluate joint ecological and evolutionary responses to heterospecific presence, we investigated whether and how population density and genotypic diversity covaried across treatments. We performed a linear model with treatment type (no-heterospecific control, or *C. striatum*, or *P. aurelia*, or *P. bursaria* treatments) and final density as explanatory variables and genotypic diversity as response variable and their interaction (*diversity* ∼ log_10_(*final_density*) * *treatment*). Temperature was excluded from the final model because it did not show a significant effect in this analysis. To facilitate interpretation, we calculated the standard deviation of the observed responses for the *T. thermophila* control, along the density and genotypic diversity axes, and used it as reference for interpreting ecological and evolutionary responses for each treatment.

Last, to understand if and how heterospecific traits influenced *T. thermophila* population density (ecological effects) and genotypic diversity (evolutionary effects), we evaluated the covariance of heterospecific growth rates and the effect size of either genotypic diversity or final density of *T. thermophila*. First, to evaluate the r-TPC of heterospecifics, we performed an ANOVA for each heterospecific with growth rate as dependent variable and temperature, as a factor, as the independent variable. We then used this data and fitted linear models of heterospecific growth rate and effect size of either population density or genotypic diversity across temperatures and heterospecific treatments. Since these data (i.e., heterospecific growth rates and effect sizes of density and genotypic diversity) were collected in different experiments, we were not able to use specific datapoints to fit a linear model. Therefore, we used linear model on bootstrapped data pooling from the distribution of the underlying data. We first generated datasets with 1000 points using the normal distributions of the growth rate data for heterospecifics at each temperature, and the *T. thermophila* effect size (final density and genotypic diversity) distributions at each temperature in the presence of each heterospecific. At each bootstrap run (n = 9999), we pooled six samples, to reflect experimental sample size, from each of these normal distributions (three heterospecific x three temperatures x six replicates = 54) and calculated the slope of the relationship using a linear model (*effect size* ∼ *growth rate*). We then generated a distribution of slopes across all temperatures and heterospecific species and calculated the mean and 95% confidence interval and performed a t-test against mean slope equal zero. All analyses were performed using R version 4.4.1 (2024-06-14).

## Results

### Intra-specific interactions and temperature affect evolution of T. thermophila

*Tetrahymena thermophila* genotypes collectively showed a 21-fold increase in per capita growth rates, from 0.054 ± 0.078 day^−1^ (mean ± SE) at 19°C, to 1.160 ± 0.042 day^−1^ at 25°C (Fig 2A, Table S2), but there was no difference in growth response across temperatures among genotypes (Table S3). These results set the expectation that genotypic frequency and diversity should remain constant across temperatures (Fig 1A), in the absence of an intraspecific interaction among genotypes when genotypes are grown together (i.e., no effect of temperature on relative genotypic frequency and diversity, gray lines in Fig 2B and 2C).

**Figure 2:**
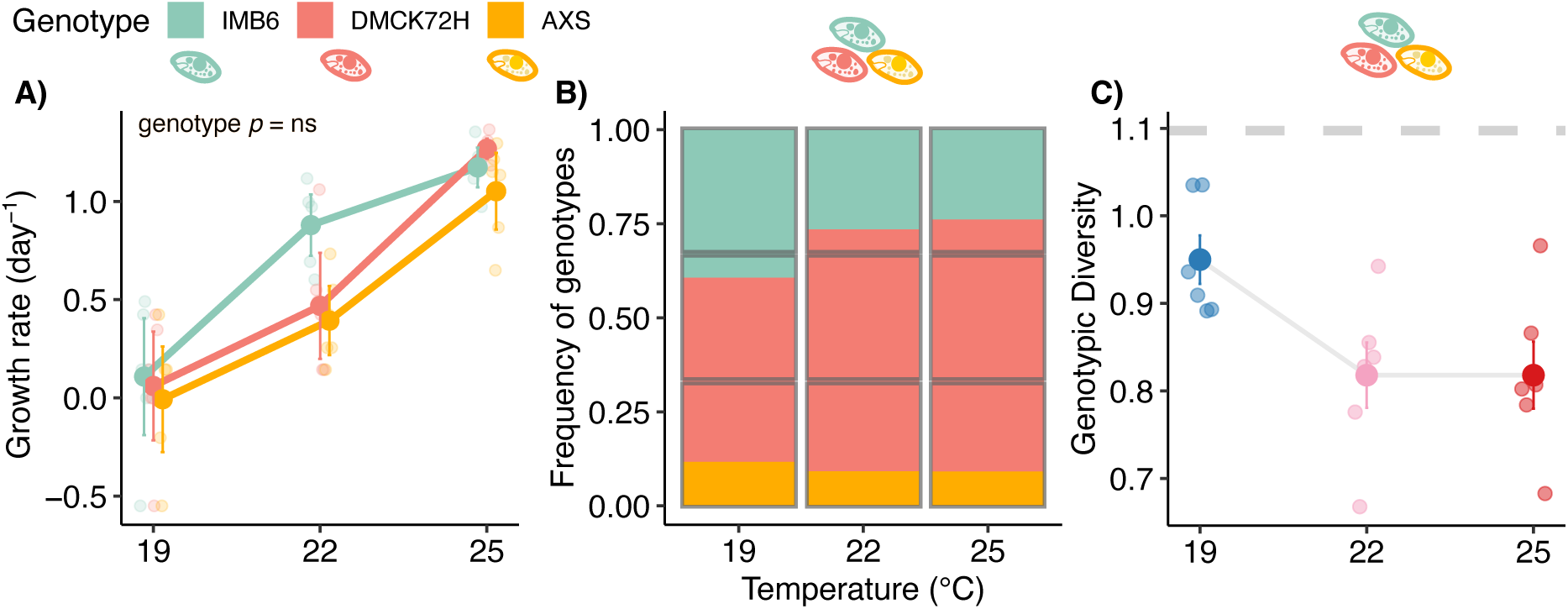
Contrasting genotype-specific intrinsic growth rates with the genotypic frequency and diversity in isolation and in co-culture across temperatures. A) Intrinsic growth rate (day^−1^) of genotypes in isolation. B) Average frequency of each genotype in co-culture. C) Genetic diversity (Shannon diversity index) of *T. thermophila* population at different temperatures. Gray boxes in B and dashed gray line in C denote expected frequencies and diversity, respectively, given that linear model showed no difference among r-TPC genotypes, as shown in A. Data shows means and standard errors. Datapoints shown in the background.

Yet, when all genotypes were grown in co-cultures, genotypic frequencies were affected by the interaction of temperature and genotype identity, with one genotype (DMCK72H, in red) becoming increasingly dominant at higher temperatures (Fig 2B, frequency ∼ temperature * genotype, *F*-value = 24.33, *p* < 0.001). This temperature-driven change in genotypic frequencies resulted in smaller genotypic diversity at higher temperatures (Fig 2C, diversity ∼ temperature, *F*-value = 6.63, *p* = 0.02), indicating that intraspecific interactions among genotypes and temperature jointly shape selection and evolution of our *T. thermophila* populations in the absence of heterospecifics.

### Inter-specific interactions and temperature affect evolution of T. thermophila

When compared to controls, the presence of heterospecifics influenced genotypic frequencies in *T. thermophila* (Fig 3A, treatment level, *F* = 4.20, *p* = 0.007), resulting in changes in genotypic diversity (Fig 3B, *F* = 6.073, *p* = 0.001). The frequency of the genotype DMCK72H increased at higher temperatures in the presence of all heterospecifics, but the magnitude to which this occurred was heterospecific-specific (Fig. 3A, temperature level, *F* = 9.263, *p* = 0.003). *Colpidium striatum* increased the dominance of DMCK72H genotype and decreased *T. thermophila* genetic diversity with increasing temperature (Fig 3B). *Paramecium aurelia* increased the dominance of the DMCK72H genotype at intermediate temperatures, but at high temperatures other genotypes increased frequency (Fig. 3A) which resulted in a U-shaped response in diversity across temperatures (Fig. 3B). Finally, in the presence of *Paramecium bursaria*, DMCK72H increased dominance, which resulted in lower diversity at higher temperatures (Fig 3B). While higher temperatures decreased genetic diversity in the no-heterospecifics controls, this decrease was much larger in the presence of *P. aurelia* and, especially, *P. bursaria* (Fig 3B, Table S4). Together, these results indicate that interspecific interactions interact with temperature to drive selection and resulting shifts in genotypic diversity in *T. thermophila*.

**Figure 3:**
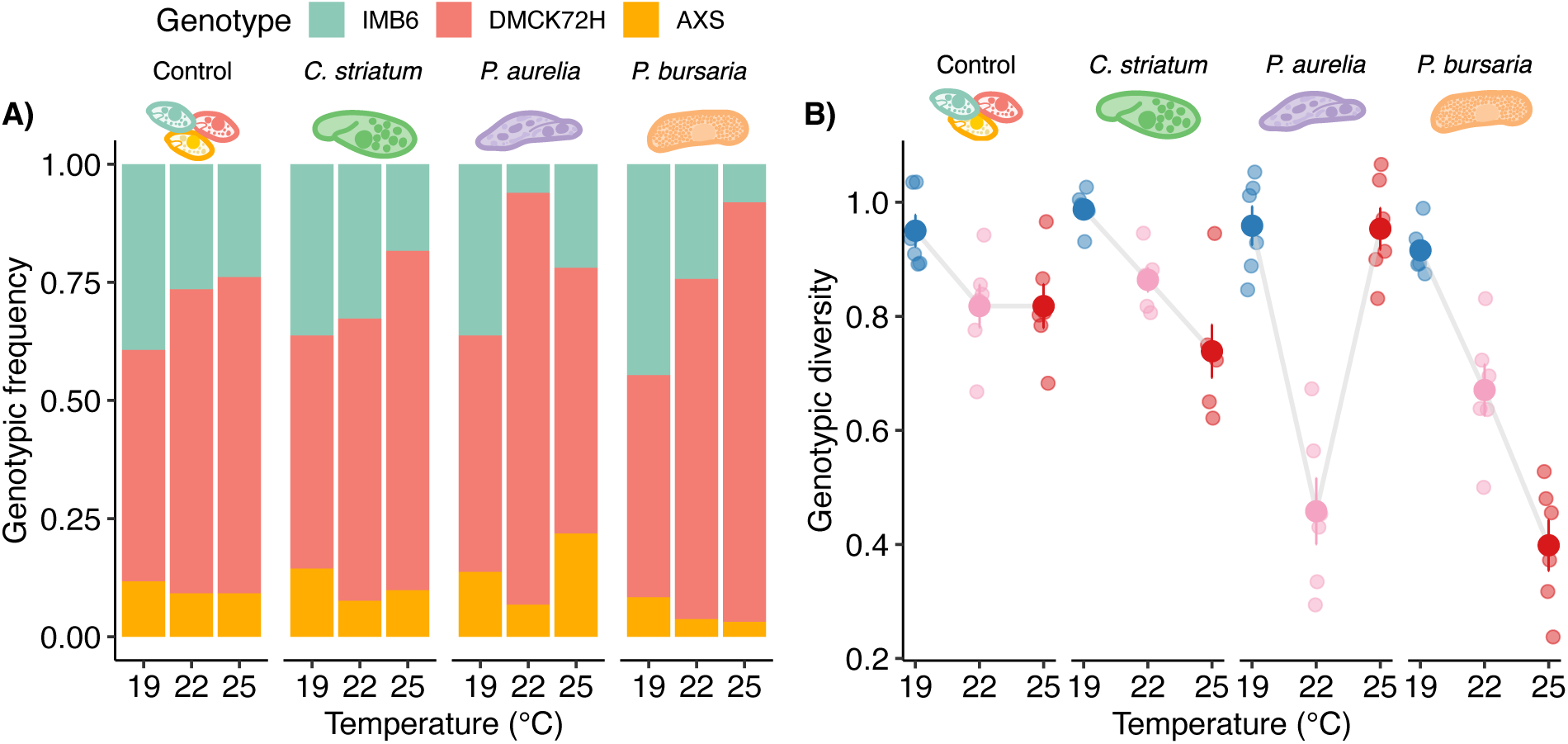
Genotypic frequency (A) and diversity (B) of *T. thermophila* in the absence (Control) and presence of heterospecific (*C. striatum*, *P. aurelia*, *P. bursaria*) across temperatures. The bars in A show average frequency. The data in B shows Shannon diversity index means and standard errors, datapoints shown in the background. The control data shown here is the same as in Fig 1B and 1C.

### Mechanisms of evolutionary response to heterospecifics

To further understand the observed variation in evolutionary responses across heterospecific treatments, we tested –and showed– that each protist had a distinct ecological effect (positive, neutral, or negative) on *T. thermophila* (i.e., population density) across temperatures (Fig 4A). Interestingly, the magnitude –and sometimes sign– of each effect changed with temperature across heterospecific treatments (Fig 4A). Specifically, *C. striatum* had an overall positive effect (at 19°C, 1.295 [1.183, 1.377], mean CI [lower, upper]) on all genotypes and population density of *T. thermophila* compared to control (Fig 4A, in green). Meanwhile, *P. aurelia* showed a negative impact on *T. thermophila* at low temperatures (–0.317 [–0.256, – 0.403]), but no effect at higher temperatures (Fig 4A, in purple). However, at intermediate temperatures, *P. aurelia* negatively impacted genotype IMB6 (–0.627 [–0.388, –1.079]), which is the driver of the decrease in diversity observed in Fig 3B. Interestingly, the effects of *P. bursaria* on *T. thermophila* density shifted from positive, at low temperatures (0.504 [0.618, 0.368]), to mostly negative at intermediate and high temperatures (–0.268 [–0.122, –0.439]) (Fig 4A, in orange). These results show that temperature and heterospecifics shape the net effect on focal species population density and magnitude of the ecological interaction with the *T. thermophila* population, and that the type of interaction in turn determines selection among genotypes.

**Figure 4:**
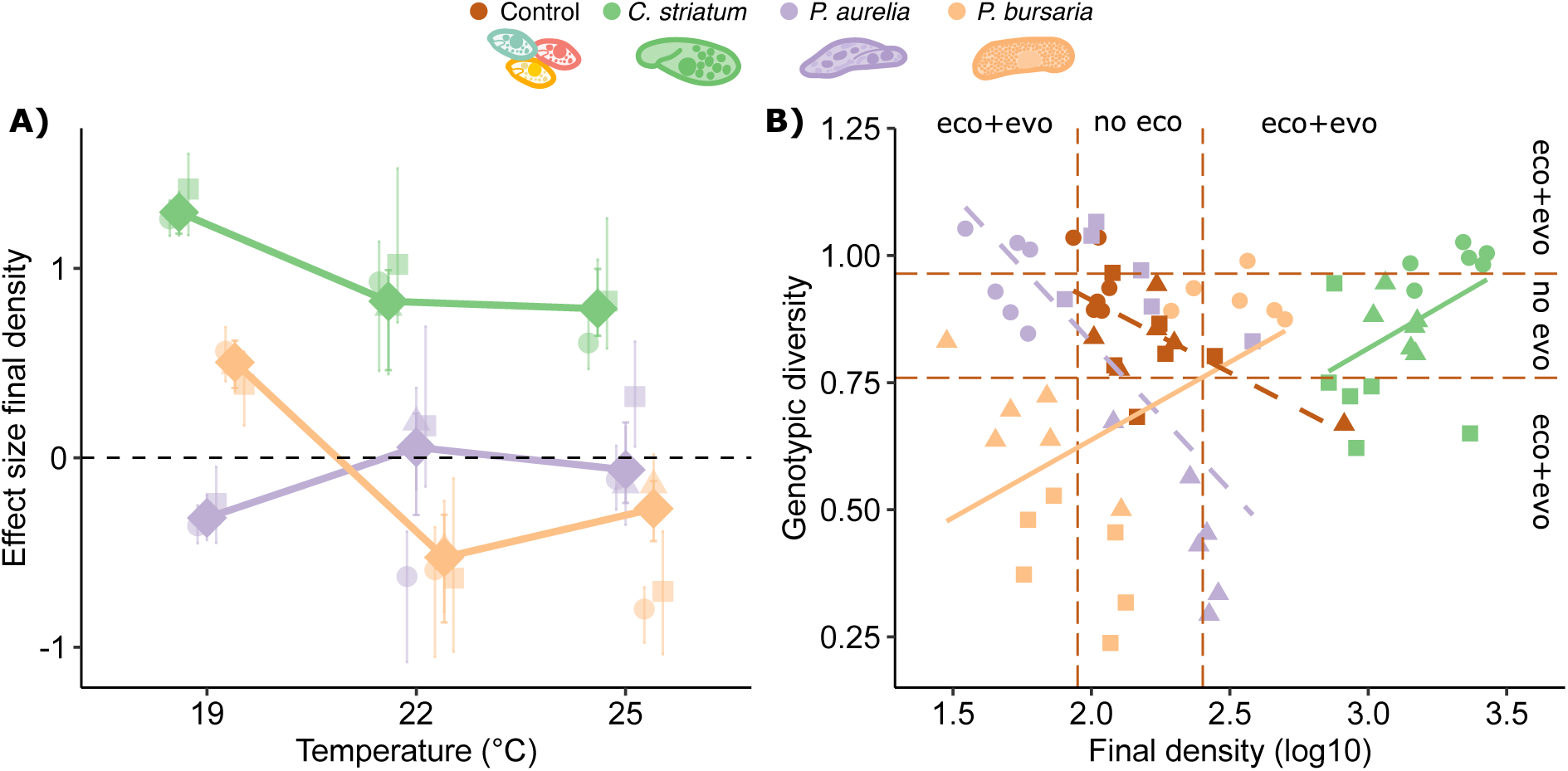
Effects of heterospecifics on *T. thermophila* population density and on the covariation between density and genotypic diversity. A) Effect size of the final population density (cell/ml) of the *T. thermophila* population (diamonds) and strain specific responses shown in the background (circles = IMB6, triangles = DMCK72H, squares = AXS). Colors represent different heterospecifics. The dashed line indicates a net zero effect size (no difference from control). B) Covariation of *T. thermophila* genotypic diversity (Shannon index) and final density of *T. thermophila* in the absence (control, brown) and presence of heterospecifics. Solid regression lines indicate slopes significantly different from the control; dashed regression lines indicate slopes not different from the control. Different symbols represent temperatures (circles = 19 °C, triangles = 22 °C, squares = 25 °C). Brown horizontal and vertical dashed lines mark the distribution (i.e., standard deviation) of the *T. thermophila* control across density and diversity, used as a reference for interpreting the magnitude of ecological and evolutionary responses relative to control. Points inside the control distributions indicate no deviation from the control baseline along that axis (“no eco” or “no evo”); points outside these distributions indicate ecological and/or evolutionary responses relative to control (“eco+evo”).

We investigated the joint ecological and evolutionary effects of heterospecific presence by analyzing the relationship between *T. thermophila* density and genotypic diversity (Fig. 4B). When analyzing the control on itself, it showed a negative correlation (slope = –0.285, t = – 3.227, p = 0.005, r^2^ = 0.394). Relative to the control (slope = –0.285, t = –1.681, p = 0.097, r^2^ = 0.394), the presence of *P. aurelia* did not significantly alter the negative covariation between density and diversity (slope = –0.583, t = –1.442, p = 0.154, r^2^ = 0.518). In contrast, both *C. striatum* (slope = 0.316, t = 2.229, p = 0.029, r^2^ = 0.211) and *P. bursaria* (slope = 0.306, t = 2.978, p = 0.004, r^2^ = 0.187) induced positive covariation between density and diversity. Deviations from the *T. thermophila* baseline control (measured as standard deviation along density and genotypic diversity axes, dashed lines, Fig 4B) show both ecological and evolutionary responses in each treatment. Specifically, a response that is larger (smaller) than the control along the density axes points to a larger (smaller) ecological response to the treatment, while a response that is larger (smaller) relative to the control along the genotypic diversity axis suggests a larger (smaller) evolutionary response to the treatment compared to control. A larger (smaller) response along both axes simultaneously suggests larger (smaller) coupled or joint ecological and evolutionary responses relative to control. These results indicate that non-neutral effects of heterospecifics (positive and negative) on *T. thermophila* density (Fig. 4A) were accompanied by corresponding shifts in the density-genotypic diversity relationship.

Finally, we investigated whether and how heterospecific’s traits influenced coupled ecological and evolutionary response of *T. thermophila*. First, we evaluated the effect of temperature on each heterospecific growth rate. Each heterospecific showed distinct r-TPC (Fig S1, Table S1) with *P. bursaria* growth rate being effectively temperature independent (*p* = 0.54) in this temperature range, *P. aurelia* displaying a similar increase with temperature compared to *T. thermophila* (*p* < 0.001), and *C. striatum* peaking at intermediate temperatures (*p* = 0.01). When we grouped these growth rate results across heterospecific species identity and temperature, we observed a positive relation between heterospecific growth rates and *T. thermophila* population density (Fig 5A, slope = 0.744, *t* = 214.51, *p* < 0.0001), and genotypic diversity (Fig 5B, slope = 0.185, *t* = 167.27, *p* < 0.0001), suggesting that heterospecific traits (i.e., growth rate) affect the ecological and evolutionary responses of *T. thermophila*.

**Figure 5:**
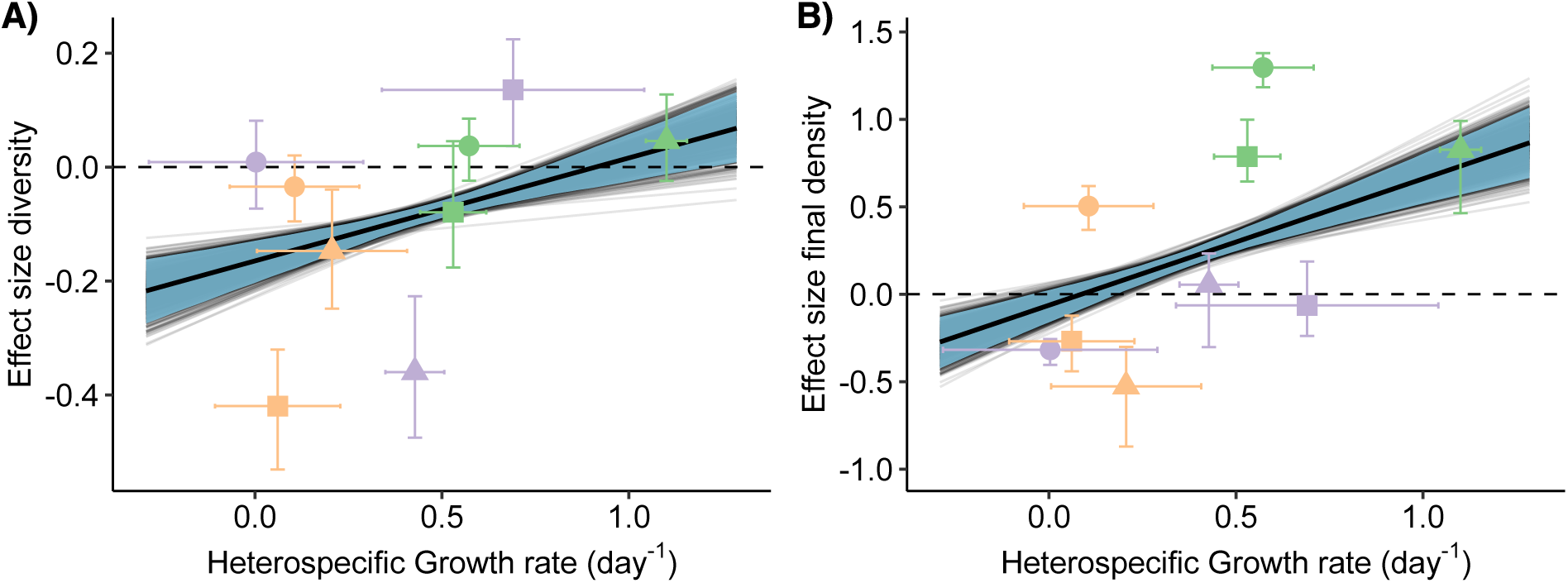
Correlation of heterospecific trait (i.e., growth rate in units of day^−1^) and *T. thermophila* final density (A) and genotypic diversity (B). Bold line represents average slope, and fine background lines each slope simulation (n = 999), of this relationship after bootstrapping both covariates, shaded area represents 95% confidence interval. Effect sizes were calculated using the mean difference between controls (without heterospecific) and treatments (with heterospecific). Shapes represent different temperatures (circles = 19 °C, triangles = 22 °C, squares = 25 °C). Colors represent different heterospecific. Error bars represent 95% confidence interval.

## Discussion

Populations can rapidly adapt to novel environmental conditions, such as elevated temperatures, through changes in additive genetic variation, often manifested as shifts in genotypic frequencies (Angilletta Jr., 2009; Barrett and Schluter, 2008; Fischer et al., 2021; Lai et al., 2019). However, biotic interactions—both within and between species—can modulate this adaptive response (Han et al., 2024; Tseng et al., 2019; Tseng and O’Connor, 2015). In this study, we show that intraspecific interactions across a temperature gradient influence evolutionary change –characterized as shifts in genotypic frequencies– in a population of the protist *Tetrahymena thermophila*, such that observed frequency change across temperatures cannot be predicted merely by the growth rate thermal response (r-TPC) of genotypes growing in isolation (Fig 2). We also found that interspecific interactions exerted strong differential selection among genotypes, which, together with temperature, determined changes in population genotypic composition by increasing or decreasing dominance of a single genotype and altering genotypic diversity (Fig 3). These effects of interspecific ecological interactions, operationally defined here as the net effect of heterospecifics on the focal population density (positive, negative, or neutral, Fig 4), in turn seems to be linked to heterospecific growth rates (Fig 5). Our results therefore highlight how evolutionary responses are affected by biotic interactions and temperature, and demonstrate the importance of investigating genotype-specific responses in the broader biotic context in which they occur to better understand rapid evolutionary responses in a changing world.

Intraspecific interactions, rather than genotype-specific r-TPCs, determined patterns of genotypic dominance across temperatures (Fig 2B). Indeed, intraspecific interactions can alter the phenotypic response of genotypes (Fridley et al., 2007), resulting in changes in the strength of the interaction, which in some cases even weaken intraspecific competition (Noto and Hughes, 2020). Interestingly, a previous study that predicted genotypic composition using r-TPC of only two genotypes of *T. thermophila* was successful in matching observed genotypic frequency of the population (Liu et al., 2024). Our results could indicate that when a larger number of genotypes are present, evolutionary outcomes deviate from predictions based on pairwise r-TPCs alone, perhaps because of higher-order interactions among multiple genotypes, that can produce non-additivity or functional change which cannot be captured by pairwise expectations (Billick and Case, 1994). Last, the genotypic sorting that results in rapid evolutionary change across treatments, can result from multiple processes, including differences in growth rates, differences in competitive traits, frequency dependence, or even some form of genotype-genotype facilitative effect. While our experimental design does not allow us to determine which of these processes more likely explain the observed pattern of evolutionary response, we can rule out it being solely based on differential growth rates across temperatures (Fig 2A), thus clearly indicating that intraspecific interactions modulate the relation between warming and selection at the population level.

Genotypic diversity of *T. thermophila* declined with increasing temperature, but this trajectory was modified by the presence of heterospecifics, even though the same genotype consistently remained dominant (Fig 3A). Both *C. striatum* and *P. bursaria* reduced the genotypic diversity of the focal population with increasing temperature, but genotypic diversity showed a U-shaped thermal response in the presence of *P. aurelia* (Fig 3B). The reduction of genotypic diversity with increasing temperature is an expected effect of climate change, which can result from selection in favor of a given genotype, or genotypic loss (Pauls et al., 2013) and can have several consequences for population and community dynamics (Hughes et al., 2008). More genotypes can confer positive, but limited, effects on population response to disturbances like increased temperatures (Singleton et al., 2021), or grazing (Hughes and Stachowicz, 2004). At the community level, increased genetic diversity within species can promote species diversity (Booth and Grime, 2003; Uriarte and Menge, 2018; Vellend, 2006) and connectivity, resulting in food web structural changes (Gibert and DeLong, 2017). Moreover, higher genotypic diversity has been shown to weaken intraspecific competition (Noto and Hughes, 2020) –an important mechanism of coexistence (Chesson, 2000)– and weaken predation, which often promotes species persistence (Gibert et al., 2015, 2015; Gibert and Brassil, 2014). Precisely investigating the effects of intraspecific diversity in rapidly shifting biotic and abiotic contexts is thus a necessary step towards understanding eco-evolutionary outcomes in rapidly warming communities.

Interspecific interactions had both positive and negative effects on focal population density (Fig 4A). Concomitantly, these interactions influenced evolutionary trajectories, and while these responses were temperature-dependent and species-specific, they were determined by the traits of heterospecifics (Fig 5). For example, *C. striatum* facilitated *T. thermophila* populations across temperatures, while *P. aurelium* had a weakly competitive effect at low temperatures which became neutral at warmer temperatures. *Paramecium bursaria*, in contrast, shifted from facilitation at low temperatures to competition at high temperatures. Changes from competition to facilitation can occur along a temperature gradient (Olsen et al., 2016), and as a result of changing resources (Alba et al., 2019). This shift in the direction of ecological interactions is consistent with patterns described by the stress gradient hypothesis (SGH, Bertness and Callaway, 1994), which predicts context-dependent shifts between facilitation and competition. For example, we found a positive correlation between genotypic diversity and population density only when heterospecifics either facilitated (*C. striatum*) or inhibited (*P. bursaria*) *T. thermophila* population density (Fig 4B). In contrast, when the heterospecific effect on the focal species was neutral (e.g., *P. aurelia*; Fig. 4A), the relation between genotypic diversity and population density was not different from the controls (Fig. 4B). Importantly, however, temperatures were not stressful enough to the focal species for our results to be fully interpretable within the boundaries of the SGH *sensu stricto*, even if our results align with its expectations. We hope that understanding why that is may be the subject of future studies. Together, these results demonstrate that temperature sets the ecological context (i.e., competition, facilitation) in which evolution happens, while ecological interactions mediate evolutionary responses to temperature by shaping genotypic diversity, with reciprocal feedbacks between ecological and evolutionary dynamics.

The dominant genotype-specific responses to heterospecifics conferred population-level benefits by reducing the impact of interspecific competition (Fig 4A). For example, while subordinate genotypes were negatively impacted by *P. bursaria* at high temperatures, the dominant genotype “rescued” the *T. thermophila* population at 25° C (Fig 4A, orange triangle). Genotype-specific competitive ability, documented in plants (Johnson et al., 2008), can shift under changing environmental conditions (Fridley et al., 2007) and with genotypic differences in both intra- and interspecific interactions that also interact with warming (Taylor et al., 2019). These results suggest that larger standing genetic variation can provide complimentary effects that can promote population performance across ecological contexts. Thus, genotype-specific interspecific competitive abilities shape population resilience under biotic and abiotic stress, emphasizing the role of genetic variation in mediating eco-evolutionary responses to environmental change.

Our results suggest that variation in heterospecific growth rates, rather than species identity alone, underlies the ecological and evolutionary effects we observed (Fig 5). Indeed, growth rate is a key functional trait that can directly affect ecological interactions, especially in protists which is correlated with body size and intake rates (Weisse et al., 2016; Wieczynski et al., 2021). Larger body size increases interaction strength of consumption in protists (DeLong and Vasseur, 2012), which might lead to stronger negative effects through exploitation (Tilman, 1980). Here, all heterospecifics are larger than the focal species (Wieczynski et al., 2021), yet we observed positive ecological effects, which might be due to differential predation patterns on bacteria. For example, a recent study showed that *Colpidium* sp. and *Tetrahymena pyriformis* alters the bacterial community composition differently, therefore suggesting that they do not in fact eat the same bacterial species (Rocca et al., 2022), despite long-standing assumptions to the contrary (Fox and Morin, 2001; Jiang and Morin, 2007). In our experiment, when the two protists are together, *Colpidium* effects on bacterial community might facilitate fractions of the bacteria consortia that are mostly exploited by *Tetrahymena*. However, because bacterial community was not measured, the extent to which these positive effects are driven by indirect, bacteria-mediated mechanisms, remains unresolved. Nonetheless, while our experimental design and results do not target specific traits responsible for this relationship, our findings suggest that heterospecifics’ key traits, such as intrinsic growth rates, may shape evolutionary responses of focal species across environmental gradients.

Overall, our findings highlight that evolutionary responses to environmental change cannot be predicted by abiotic factors alone. Rather than yielding general predictions, our results show that evolutionary responses to temperature are contingent on biotic context, specifically conspecific and heterospecific interactions, therefore refining our current understanding in exciting new ways. Intraspecific interactions significantly altered genotypic frequencies and masked expected thermal responses, while interspecific interactions further shaped ecological and evolutionary outcomes in species- and temperature-dependent ways. The dominance of a single genotype under varying biotic and abiotic contexts suggests strong selection, potentially limiting genotypic diversity and adaptive potential at higher temperatures. Moreover, the presence of positive interactions under stress underscores the importance of considering both competitive (i.e., negative) and positive interactions in eco-evolutionary models. These results emphasize the need to integrate biotic context into predictions of evolutionary dynamics under global change scenarios.

## STATEMENT OF AUTHORSHIP

MHL, JPG designed the study. MHL, AY, ZH, KD collected the data. EL and MHL conducted data analyses. EL wrote the first draft of this manuscript with contribution of MHL. All authors contributed significantly to revisions.

## ACKNOWLEDGMENTS

This work was supported by NSF DEB award number 2224819, NSF CAREER award number 2337107, and a Simons Foundation Early Career Fellowship in Aquatic Microbial Ecology and Evolution number LS-ECIAMEE-00001588 to JPG.

