## Supplemental_material for "Ecological interactions mediate evolutionary responses to temperature in microbial communities"

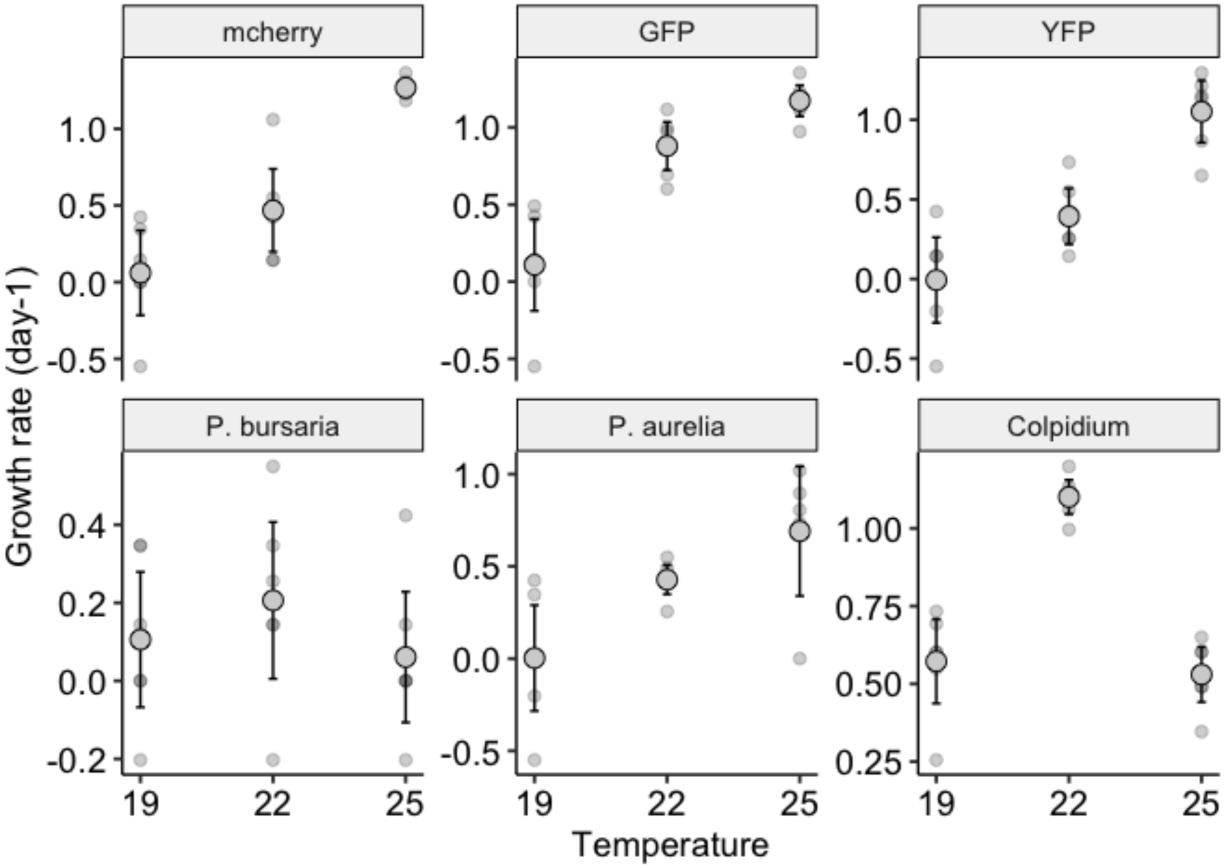


Figure S1: Growth rate response at different temperatures for Tetrahymena strains (DMCK72H, IMB6, AXS) and protist heterospecific (Paramecium bursaria, Paramecium aurelia, Colpidium). Data represents means ± confidence interval (95%).

Tables

Table S1: *Tetrahymena termophila* genotypes provenance.

| Name | Original Provenance | Mutations/Notes |
|---|---|---|
| AXS (YFP) | Chalker Lab, Washington University in St. Louis | YFP fusion into the BTU1 locus of Tetrahymena Strain CU428.2 (RRID TSC SD00178) |

| | |
|---|---|
| DMCK72H (mCherry) | Chalker Lab, Washington University in St. Louis |
| IMB6 (GFP) | Chalker Lab, Washington University in St. Louis |

Table S2: Statistical results for the effects of temperature on the individual growth rate of the T. thermophila strains, and heterospecifics used in the experiment, using ANOVA with temperature as independent categorical variable and growth rate as dependent variable.

| Organism | | Sum Sq | Df | F value | Pr(>F) |
|---|---|---|---|---|---|
| *Tetrahymena* strains | | | | | |
| DMCK72H | | | | | |
| | Temperature | 4.53 | 2 | 28.53 | p<0.001 ** |
| | Residuals | 1.19 | 15 | NA | NA |
| IMB6 | | | | | |
| | Temperature | 3.63 | 2 | 28.26 | p<0.001 ** |
| | Residuals | 0.96 | 15 | NA | NA |
| AXS | | | | | |
| | Temperature | 3.43 | 2 | 23.33 | p<0.001 ** |
| | Residuals | 1.1 | 15 | NA | NA |
| Protist heterospecific | | | | | |
| *P. bursaria* | | | | | |
| | Temperature | 0.07 | 2 | 0.65 | 0.54 |
| | Residuals | 0.77 | 15 | NA | NA |
| *P. aurelia* | | | | | |
| | Temperature | 1.34 | 2 | 7.03 | 0.01 ** |
| | Residuals | 1.33 | 14 | NA | NA |
| *Colpidium* | | | | | |
| | Temperature | 1.21 | 2 | 39.86 | p<0.001 ** |

| | | | | | |
|---|---|---|---|---|---|
| | Residuals | 0.23 | 15 | NA | NA |

Table S3: Summary statistics for effects of temperature on growth rate of Tetrahymena strains using a linear model with temperature as covariant, strain as factor and growth rate as independent variable. (growth ~ temperature * strain).

| | Sum Sq | Df | F value | Pr(>F) |
|---|---|---|---|---|
| (Intercept) | 0.02 | 1 | 0.31 | 0.58 |
| Strain | 0.04 | 2 | 0.28 | 0.76 |
| Temp | 4.53 | 2 | 31.29 | 0 |
| Strain:Temp | 0.48 | 4 | 1.66 | 0.18 |
| Residuals | 3.26 | 45 | NA | NA |

Table S4: Statistical results for the frequency of the T. thermophila strains across temperature and presence of other protists (frequency ~ temperature * strain * heterospecific).

| | Sum Sq | Df | F value | Pr(>F) |
|---|---|---|---|---|
| (Intercept) | 0.165 | 1 | 21.381 | 0.000 |
| temp | 0.071 | 1 | 9.263 | 0.003 |
| strain | 0.100 | 2 | 6.509 | 0.002 |
| heterospecific | 0.097 | 3 | 4.200 | 0.007 |
| temp:strain | 0.170 | 2 | 11.034 | 0.000 |
| temp: heterospecific | 0.098 | 3 | 4.252 | 0.006 |
| strain: heterospecific | 0.302 | 6 | 6.523 | 0.000 |
| temp:strain: heterospecific | 0.331 | 6 | 7.147 | 0.000 |
| Residuals | 1.480 | 192 | NA | NA |

| Estimate | Std. Error | t value | F-value | |
|---|---|---|---|---|
| (Intercept) | 0.8647556 | 0.1870158 | 4.6239703 | 0.8179733 |
| temp | -0.0257130 | 0.0084485 | -3.0434931 | 0.9848322 |
| strain DMCK72H | -0.9229598 | 0.2644803 | -3.4897107 | 0.6144499 |
| strain AXS | -0.6713071 | 0.2644803 | -2.5382119 | 0.8697607 |
| trtColp | 0.0826284 | 0.2644803 | 0.3124179 | 0.0000000 |
| trtPaur | -0.1242612 | 0.2644803 | -0.4698314 | 0.8013897 |
| trtPbur | 0.7336009 | 0.2644803 | 2.7737449 | 0.1330712 |
| temp:strainDMCK72H | 0.0556495 | 0.0119480 | 4.6576399 | 0.5081724 |
| temp:strain AXS | 0.0214895 | 0.0119480 | 1.7985840 | 0.9571172 |
| temp:trtColp | -0.0041193 | 0.0119480 | -0.3447677 | 0.0000000 |
| temp:trtPaur | 0.0017933 | 0.0119480 | 0.1500900 | 0.8718495 |
| temp:trtPbur | -0.0352639 | 0.0119480 | -2.9514460 | 0.1337395 |
| strain DMCK72H:trtColp | -0.2463330 | 0.3740317 | -0.6585886 | 0.2150431 |
| strainYFP:trtColp | -0.0015522 | 0.3740317 | -0.0041499 | 0.0015422 |
| strain DMCK72H:trtPaur | 0.5995917 | 0.3740317 | 1.6030506 | 0.9067635 |
| strain AXS:trtPaur | -0.2268082 | 0.3740317 | -0.6063876 | 0.9746499 |
| strain DMCK72H:trtPbur | -1.5150692 | 0.3740317 | -4.0506443 | 0.9465051 |
| strain AXS:trtPbur | -0.6857336 | 0.3740317 | -1.8333571 | 0.3887024 |
| temp:strain DMCK72H:trtColp | 0.0116516 | 0.0168970 | 0.6895628 | 0.0487585 |
| temp:strainYFP:trtColp | 0.0007063 | 0.0168970 | 0.0418000 | 0.0060113 |
| temp:strain DMCK72H:trtPaur | -0.0214083 | 0.0168970 | -1.2669847 | 0.9564118 |
| temp:strain AXS:trtPaur | 0.0160285 | 0.0168970 | 0.9485958 | 0.9677753 |
| temp:strain DMCK72H:trtPbur | 0.0749541 | 0.0168970 | 4.4359331 | 0.9715647 |

| Estimate | Std. Error | t value | F-value | |
|---|---|---|---|---|
| temp:strain AXS:trtPbur | 0.0308376 | 0.0168970 | 1.8250294 | 0.3933806 |

**Table S5**: Statistical results for the genotypic diversity index of the T. thermophila population across temperature and presence of other protists (diversity ~ temperature * heterospecific).

| | Estimate | Std. Error | t value | Pr(>\|t\|) |
|---|---|---|---|---|
| (Intercept) | 1.3463244 | 0.3265936 | 4.1223226 | 0.0001100 |
| trtColp | 0.4283522 | 0.4618732 | 0.9274239 | 0.3571909 |
| trtPaur | -0.5365638 | 0.4618732 | -1.1617123 | 0.2496666 |
| trtPbur | 1.2114790 | 0.4618732 | 2.6229690 | 0.0108832 |
| temp | -0.0220172 | 0.0147540 | -1.4922851 | 0.1405377 |
| trtColp:temp | -0.0194079 | 0.0208653 | -0.9301523 | 0.3557875 |
| trtPaur:temp | 0.0211267 | 0.0208653 | 1.0125298 | 0.3150975 |
| trtPbur:temp | -0.0641718 | 0.0208653 | -3.0755247 | 0.0030892 |

| | Sum Sq | Df | F value | Pr(>F) |
|---|---|---|---|---|
| (Intercept) | 0.400 | 1 | 16.994 | 0.000 |
| heterospecific | 0.360 | 3 | 5.109 | 0.003 |
| temp | 0.052 | 1 | 2.227 | 0.141 |
| heterospecific:temp | 0.428 | 3 | 6.073 | 0.001 |
| Residuals | 1.505 | 64 | NA | NA |
